# Family variation in surface feeding behavior of steelhead fry predicts growth rate under hatchery conditions

**DOI:** 10.1101/2020.08.07.241141

**Authors:** Madeleine C. Wrey, James C. Skaar, Stanley B. Cates, Stephanie R. Bollmann, Claudio Fuentes, Michael S. Blouin

## Abstract

Hatchery reared salmonid fish often have lower fitness than their natural-origin counterparts when spawning in the wild. Although this difference appears to result from rapid adaptation to captivity, it is not known what traits are under selection. We hypothesize that variation in traits that confer a growth rate advantage to some individuals in the novel hatchery environment are under strong selection because survival at sea is correlated with size at release. Here we show that full sibling families of steelhead trout (*Oncorhynchus mykiss*) show substantial variation in propensity to feed at the surface as fry, and that the more surface-oriented families grew faster under hatchery conditions. We hypothesize that surface-oriented fry gain an initial growth advantage that persists through size at release. Because surface orientation is a correlate of generalized boldness, hatcheries may inadvertently select for that phenotype, which could explain the fitness differences observed between hatchery and natural-origin fish in the wild.

## Introduction

Hatchery-reared salmonid fish generally have lower fitness than natural-origin fish when spawning in the wild, and this effect is true even for early-generation hatchery fish (1–7). Common garden experiments in steelhead show that the fitness reduction appears to be genetic, rather than simply an environmental effect of the hatchery (1,6,8). That first-generation hatchery steelhead perform worse than natural-origin fish in the wild, but much better as broodstock in the hatchery, suggests that the genetic difference results from rapid adaptation to captivity (9). Unfortunately, it is not known what traits are under selection in hatcheries, nor the environmental conditions that impose the selection. Understanding those selective pressures might allow hatchery managers to modify hatchery conditions to reduce the rate of domestication, thereby improving the fitness of hatchery fish in the wild. Therefore, we are interested in identifying traits that might be under strong selection in salmon hatcheries.

The size of hatchery salmon at release is positively correlated with their probability of survival at sea (10–15). Therefore, a plausible hypothesis to explain rapid domestication is that hatcheries select for physiological or behavioral traits that allow some fish to grow quickly in the unnatural conditions in a hatchery (11,16,17). Such selection might be especially intense on steelhead, which are typically raised to smolting in one year in hatcheries versus the normal two years that they take in the wild (18,19). If those hatchery-favored traits are maladaptive in the wild, then that could explain why hatchery fish quickly evolve to have lower reproductive success than natural-origin fish in the wild environment.

Reduced anxiety/fearfulness is an almost universal outcome of domestication in vertebrates (20). Traits such as quickness to feed, and propensity to be near the surface in fish can all be manifestations of variation in general fearfulness/anxiety, which has been well characterized in fish as the shyness/boldness behavioral syndrome (21). Bolder individuals of many taxa tend to feed more readily (22), and thus it has been suggested that the high-food, high-density, predator-free environment of a hatchery selects for generalized boldness (23–26). Such selection could then explain the reduced reproductive success of hatchery fish in the wild, as the boldness that served them well in captivity could be a liability for their offspring in the low-food, predator-filled wild environment (27–29). Several studies showed domesticated salmon stocks feed more readily, and are more susceptible to, or less fearful of, predators than natural-origin stocks(23,27,30–32). Hatchery masu salmon also showed greater propensity to feed near the surface (33), in addition to reduced fear of predators and greater feeding rates (34) than natural-origin stocks. Such behavioral change is not limited to salmonids. Domesticated zebrafish showed a higher degree of surface orientation, a reduced startle response, and higher growth rate in the lab than wild zebrafish (35). It was also shown that zebrafish selected for swimming at the front of the tank (nearest to the observer) also swam higher in the water column, and fed more quickly as a correlated response to selection (36). So there appears to be a suite of genetically correlated traits in fish, including surface orientation, that all change together in response to selection for any one of them.

Steelhead fry in Oregon hatcheries are usually trained to feed on artificial food in Canadian troughs (round-bottom metal troughs of about a meter in width and depth) for several weeks before they are moved to concrete raceways to be reared through smolting. During this training period, finely-ground feed is sprinkled by hand on the water surface. Anecdotal observations from hatcheries that use natural-origin fish as broodstock suggest that some fry quickly learn to rise to the surface for food, while others cower at the bottom and fail to thrive. We hypothesize that this variation reflects heritable variation along the shyness-boldness axis, and that the bolder families get an initial growth advantage that they maintain through the rest of the rearing period. If these bolder families wind up larger at release, then subsequent size-selective mortality at sea would result in selection for boldness. If it is indeed true that bolder fish gain an initial growth advantage during the first few days of learning to feed on artificial food, then it may be possible to reduce their initial advantage by altering the feeding method. In particular, if shy fish are startled by humans, then using automated feeders might reduce the performance difference between bold and shy families. Here we tested whether families that scored higher on a correlate of boldness (propensity to feed at the surface as fry) also grew faster in mixed-family hatchery tanks, and whether that relationship was stronger when fish were hand-fed versus fed by automatic feeders.

We used a group of 11 full-sibling families of steelhead to test the following predictions: (1) Steelhead families vary substantially in propensity to feed at the surface during the critical learning period. (2) Families having a higher surface feeding score (presumably the bolder families) would also grow the fastest under hatchery rearing conditions. (3) The difference between surface-feeding and non-surface feeding families in growth would be greater when fish are fed by hand than when they are fed by automated feeders.

## Methods

### Spawning and husbandry

Winter-run steelhead were collected from the Wilson River, Oregon, by staff at the Trask Hatchery (Oregon Department of Fish and Wildlife). For our experiments we used eleven full-sibling families created using first-generation hatchery fish as broodstock (i.e. they had natural-origin parents). Spawning occurred on March 20^th^ (3 families) and on March 29^th^, 2018 (8 families). After fertilization and water hardening, embryos from each cross were transported to the Oregon Hatchery Research Center in Alsea, Oregon (OHRC).

To ensure that fry from all families would be ready for first exogenous feeding at the same time, embryos from the three families spawned on March 20^th^ were maintained on chilled water at the OHRC until the remaining families arrived. Eggs from each family were reared in individual, plastic isolates in Heath trays until all fry had hatched and were ready to begin exogenous feeding.

Prior to first feeding, fry were transported to Oregon State University’s Fryer Aquatic Animal Health Laboratory in Corvallis, OR. At the Fryer lab, fish from each family were assayed for their propensity to feed at the surface during the first 7 days of exogenous feeding. Their siblings were raised in mixed-family groups in 400 L outdoor tanks for 32 weeks in order to measure each family’s growth. These 400 L tanks were fed either by hand or using automatic feeders in order to determine if any relationship between family-average propensity to feed at the surface and family-average growth could be altered by feeding method. Fish in all experiments were fed standard hatchery diets produced by Bio-Oregon (Longview, WA) according to manufacturer recommendations based on water temperature and average body size. These recommendations ensured that an excess of food was available daily. Samples of fish in the 400 L tanks were measured biweekly to track growth and adjust feed amounts. All fish were raised and handled in accordance with Oregon State University’s Institutional Animal Care and Use Committee guidelines (Animal Care and Use Protocol # 5060).

### Quantifying family propensity to feed at the surface

Fifteen siblings from each family went into each of three, replicate, 25L flow-through indoor plastic buckets with lids (i.e. 15 siblings per bucket, three buckets per family). Inflow was positioned below water surface and aerators were turned off prior to each feeding to ensure that surface disturbance was minimized during feeding. Each bucket received 3% biomass in feed hand-sprinkled on the surface, divided equally between 10 daily feedings. Propensity for surface orientation in response to feed was quantified as the number of times the surface was broken by any fish in the bucket in response to feed during the first 30 secs after feed dispersal (hereafter number of “nose-pokes”). Number of nose-pokes was recorded daily during the first feed of the day from 30 May 2018 to 6 June 2018. The final score for each bucket was the total number of nose pokes recorded in that bucket during that seven-day period. We averaged the scores from the three buckets per family to assign a single surface-feeding score to each family. We tested the family effect on variation among buckets using a one-way ANOVA, and quantified the variation among families as the intra-class correlation (ICCest in R, version 2.15.1).

### Quantifying family Growth Rate

In order to quantify the growth of each family under hatchery conditions, an equal number of offspring from each family were assigned to be raised in mixed-family groups in each of nine 400 L tanks. During transport of the unfed fry from the OHRC to the Fryer lab there was a mortality event that resulted in variation in total numbers of fish per tank and unequal numbers of fish from each family going into each tank. This included some families being entirely missing from some tanks, although all families were represented in all treatments (S1 Table).

Fry were ponded directly into the 400 L tanks on the same day that their siblings were set up in 25-L buckets to quantify surface feeding (May 30^th^, 2018). Tanks were randomly assigned to one of three feeding treatments, with three replicate tanks per treatment. The first treatment was fed by hand, as this style of broadcast feeding is typical in hatchery settings. The second two treatments utilized the same amount of feed, but using belt autofeeders to allow fish to feed without interacting with humans. The first autofeeder treatment (“auto-bolus”) mimicked the hand feeding schedule with parcels of food entering the tank with each round of hand feeding. The second autofeeder treatment (“auto-trickle”) provided a continuous trickle of food throughout the day.

Fish from the 400 L tanks were sampled after 32 weeks of growth. We measured fork length of each fish and took fin clips for genotyping. Fish were assigned back to families via microsatellite genotyping using the SPAN B suite of loci (37) on an ABI 3730 capillary electrophoresis system with GeneMapper version 4.1 to call peaks (Applied Biosystems, Foster City, California). The SOLOMON program (38) was used with pure exclusion to assign each offspring back to its parental pair.

### Statistical Analysis

In initial exploratory analysis of the growth rate data we observed a very large tank effect that was mostly explained by the total number of surviving fish collected from each tank (S1 Fig). To adjust for that effect, we regressed the length of each fish on the total number of fish in its tank for the entire dataset, and then used the residuals from that regression as tank-adjusted measure of body size in the subsequent analyses (S2 Fig).

In order to test the hypothesis that families having high propensity to feed at the surface would also grow faster under hatchery conditions, we first obtained the average tank-adjusted body size for each family in each of the three treatments (i.e. three values per family, 33 data points total). Then using analysis of covariance (ANCOVA) we modeled the average tank-adjusted body size per family as the response variable, in terms of nose-pokes per family as the covariate and feeding treatment as the categorical variable, including the nose-pokes-by-treatment interaction. The interaction term tested whether the relationship between nose-pokes and family growth rate depended on feeding treatment. The ANCOVA was run using the GLM (general linear model) module in SYSTAT 13.2.

## Results

Propensity for surface orientation in each bucket (nose-pokes) differed significantly among families (Figure 1; ANOVA: F=12.04, df=10, P< 0.001). Indeed, the intra-class correlation among buckets from the same family = 0.79 (95% CI 0.5355 - 0.9299). Thus, steelhead fry show surprisingly large variation among families in their propensity to feed at the surface during the critical period when they are first learning to eat artificial food.

**Fig 1.**
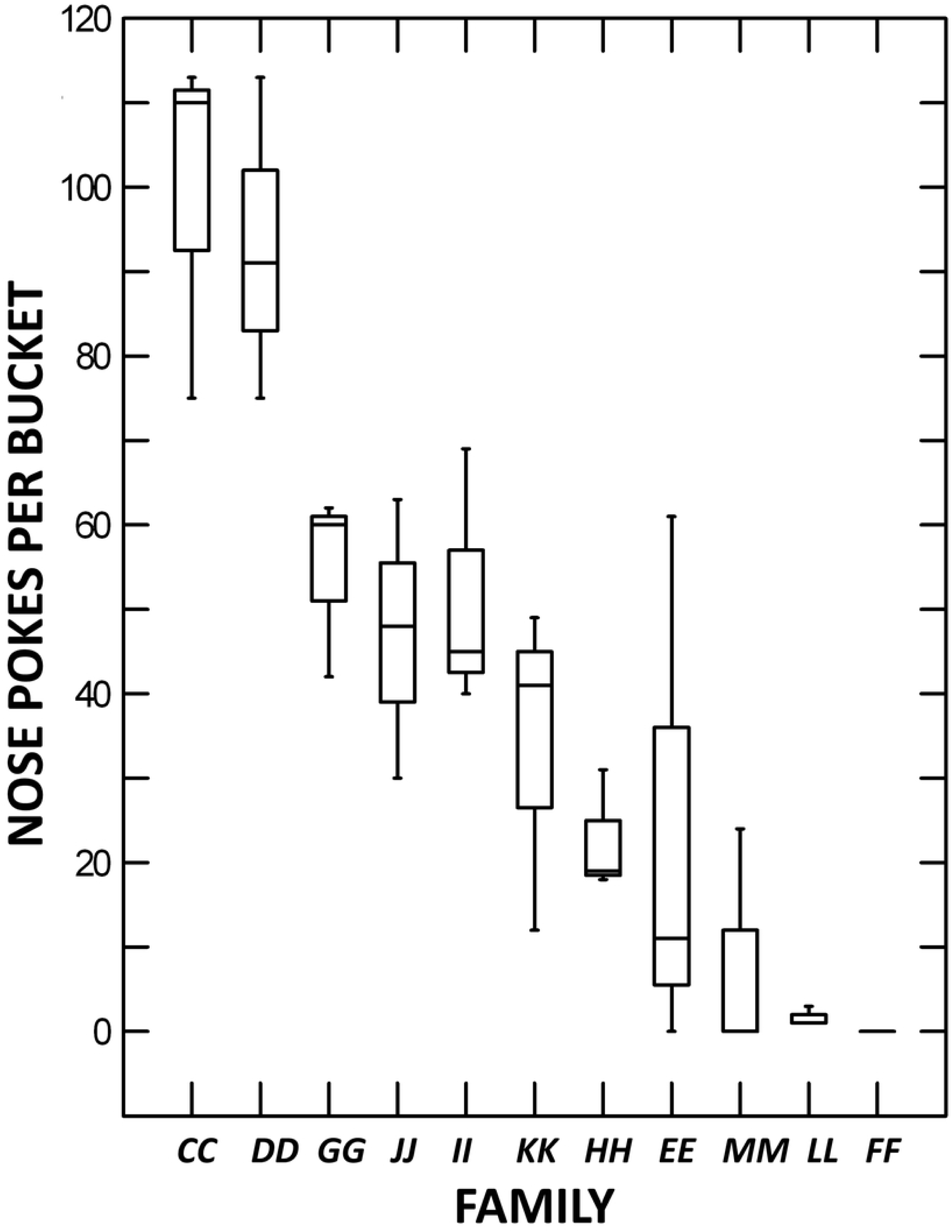
Propensity to feed at the surface by family, as measured by total nose-pokes per bucket over the first seven days of feeding. The 11 families are labelled on the X-axis. Shown are box plots for the three buckets from each family. Family identity is highly significant, explaining 79% of the variance among buckets in total nose-pokes.

The ANCOVA shows that family-average adjusted length is significantly and positively associated with family-average propensity to feed at the surface (nose-pokes) (P = 0.003; Figure 2 and Table 1). However, the nose-pokes by feeding-treatment term was not significant (Table 1), meaning that the relative performance of the different families did not depend on how they were fed. In separate regressions within each treatment group, the nominal regression coefficient was largest in the automatic-trickle treatment. So there is not even suggestive evidence that the relationship between size and nose-pokes is stronger in the hand fed treatment. There was no significant main effect of feeding treatment on the average size of fish in each family, although the average body size of families was slightly larger in the hand-fed treatment (Fig. 2).

**Table 1.** Analysis of Covariance for effects of feeding treatment and nose-pokes (covariate) on average residual length of each family.

| Source | SS | Df | Mean<br>Square | F-Ratio | P-Value |
| --- | --- | --- | --- | --- | --- |
| Treatment | 101.23 | 2 | 50.61 | 0.880 | 0.426 |
| Nose-pokes | 633.16 | 1 | 633.16 | 11.01 | 0.003 |
| Interaction | 33.91 | 2 | 16.96 | 0.295 | 0.747 |
| Error | 1,553.19 | 27 | 57.5 |  |  |

**Figure 2.**
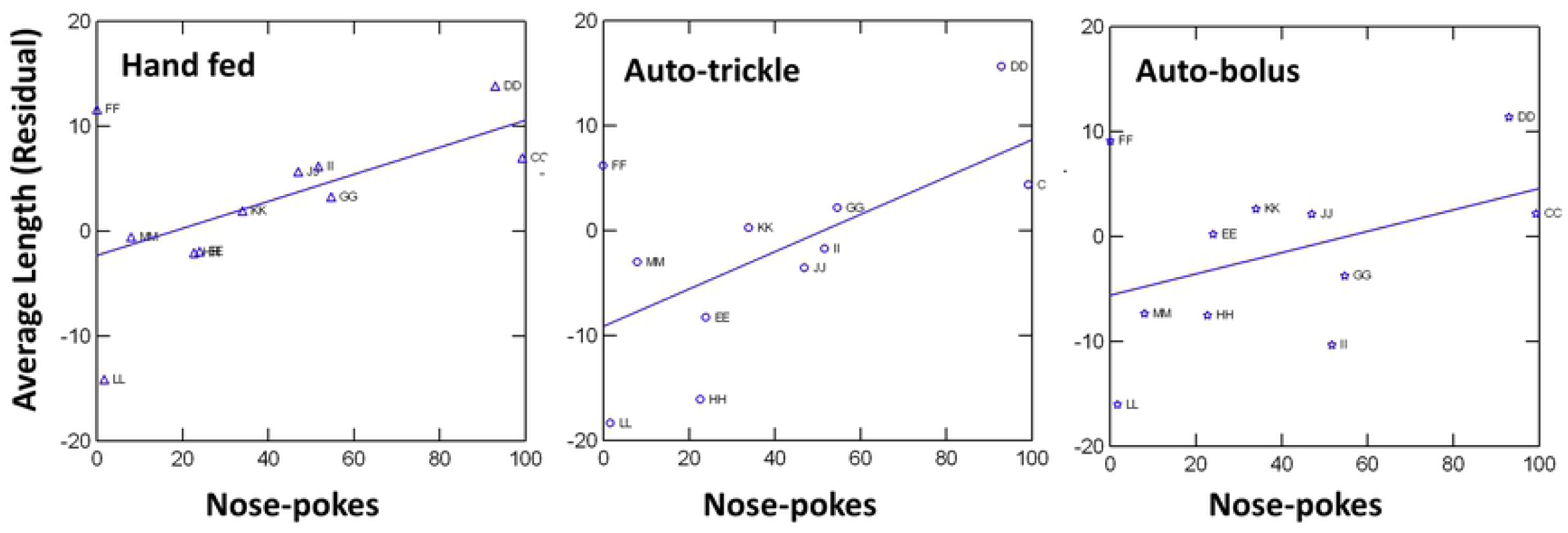
Average residual length of each family regressed against average nose-pokes (surface feeding score) of each family. Shown are least squares regression lines for each feeding treatment. The body size of each family at sampling is significantly and positively associated with its surface-feeding score. There was no main effect of treatment, nor was there a treatment by nose-pokes interaction.

One possible caveat to the result is that three of the families (CC, DD and EE) were spawned 9 days earlier than the other eight. CC and DD were among the largest, and also had the highest average nose-pokes values, although EE was unremarkable (Fig 2). As is standard hatchery practice, the three earlier families were kept in cooled water for a week to delay their development until the rest were spawned in order to ensure all families were ponded at similar sizes. Thompson et al. (39) showed, using a different steelhead hatchery program, that families from earlier spawn dates wind up larger at release, presumably owing to the cooling effect. This effect was significant in a year when spawning was spread evenly over a seven-week period, although not in a year when it was spread mostly over a two-week period. Thus, although it is conceivable that the sizes of families CC and DD were slightly inflated by the cooling, the effect of a 9-day difference is likely to be very small. We cannot statistically adjust for spawn date because there are so few families, and families CC and DD make spawn date highly confounded with nose-pokes. We therefore estimated the possible effect using the size vs. spawn-date relationships from Thompson et al. (39). If we reduce the average size of families CC, DD and EE by up to 4 mm (a greater effect than predicted by Thompson et al.’s data for fish of this size range), the results remain unchanged.

## Discussion

We found surprisingly large variation among families in their propensity to feed at the surface during their first week of exogenous feeding. Furthermore, this variation predicted which families grew fastest in the 400 L, mixed-family outdoor tanks. Given surface orientation and feeding is considered a correlate of generalized boldness, these results are consistent with the hypothesis that hatcheries inadvertently select for bolder phenotypes. The idea here is that bolder fish gain an early growth advantage that persists through size at release. Larger smolts are more likely to survive at sea, so fish from the bolder families would be over-represented in the cohort of returning adults, resulting in rapid evolution of the trait in hatchery stocks. Given that overly bold behavior can have a survival cost in the wild, selection for this trait could explain why hatchery fish produce fewer surviving offspring than do natural-origin fish when both spawn in the wild.

If hatcheries are indeed selecting for highly bold behavior in fry and juveniles, then it might be possible to alter the way fry are trained to feed in order to reduce the initial growth disparity between shy and bold fish. For example, perhaps one could separate out the initially shy fish and feed them separately until they catch up in size to the others. In this experiment, using automated feeders instead of hand-feeding did not reduce the correlation between behavior and final body size. So that change in husbandry seems unlikely to substantially reduce the intensity of selection. Of course, interventions such as this presume that a goal of the hatchery is to produce adult fish having fitness as similar as possible to that of natural-origin fish, rather than a goal of maximizing production. Regardless, these data support the hypothesis that inadvertent selection for feeding at the surface is one mechanism behind the rapid domestication of hatchery steelhead, and points to possible ways to minimize the rate of domestication.

## Acknowledgements

Thanks to the Oregon Department of Fisheries and Wildlife (ODFW), to Jen Krajcik, Joseph O’Neill and staff at the Oregon Hatchery Research Center, to Ruth Milston Clements and staff at OSU’s Aquatic Animal Heath Lab, and to the Trask hatchery for invaluable assistance in obtaining eggs and raising fish for these experiments. Genotyping was done at OSU’s Center for Genome Research and Biocomputing. This research was supported by grants to MB from the ODFW and Bonneville Power Administration.

---

**S1 Table. Number of fish per family in each tank at the end of the grow-out period**.

**S1 Fig. Average length of fish in each of the nine grow-out tanks at final sampling** (AVGLENGTH), plotted against the total number of surviving fish in each tank (NTOTAL). Case labels are the tank identification labels and the feeding treatment applied to each tank. The linear regression is shown.

**S2 Figure. Residuals from a regression of the length of each fish against the number of fish in its tank (NTOTAL)**. We used these residuals as our measure of body size on each fish for all subsequent analyses. Results using standardized residuals were almost identical, so we report results using these untransformed residuals because they are easier to interpret in terms of variation among families in mean body size.

